# The dynamic landscape of parasitaemia dependent intestinal microbiota shifting at species level and the correlated gut transcriptome during Plasmodium *yoelii* infection

**DOI:** 10.1101/2020.12.17.423374

**Authors:** Zong Yawen, Cheng Lei, Cheng Xiangyun, Liao Binyou, Ye Xingchen, Liu Taiping, Li Jiyao, Zhou Xuedong, Xu Wenyue, Ren Biao

## Abstract

**Background:** Malaria, caused by *Plasmodium*, is a global life-threatening infection disease especially during the COVID-19 pandemic. However, it is still unclear about the dynamic change and the interactions between intestinal microbiota and host immunity. Here, we investigated the change of intestinal microbiome and transcriptome during the whole *Plasmodium* infection process in mice to analyze the dynamic landscape of parasitaemia dependent intestinal microbiota shifting and related to host immunity.

**Results:** There were significant parasitaemia dependent changes of intestinal microbiota and transcriptome, and the microbiota was significantly correlated to the intestinal immunity. We found that (i) the diversity and composition of the intestinal microbiota represented a significant correlation along with the *Plasmodium* infection in family, genus and species level; (ii) the up-regulated genes from the intestinal transcriptome were mainly enriched in immune cell differentiation pathways along with the malaria development, particularly, naive CD4+ T cells differentiation; (iii) the abundance of the parasitaemia phase-specific microbiota represented a high correlation with the phase-specific immune cells development, particularly, Th1 cell with family *Bacteroidales* BS11 gut group, genera *Prevotella* 9, *Ruminococcaceae* UCG 008, *Moryella* and specie *Sutterella\**, Th2 cell with specie *Sutterella\**, Th17 cell with family *Peptococcaceae*, genus *Lachnospiraceae* FCS020 group and spices *Ruminococcus* 1*, *Ruminococcus* UGG 014* and *Eubacterium plexicaudatum* ASF492, Tfh and B cell with genera *Moryella* and species *Erysipelotrichaceae bacterium canine oral taxon* 255.

**Conclusion:** There was a remarkable dynamic landscape of the parasitaemia dependent shifting of intestinal microbiota and immunity, and a notable correlation between the abundance of intestinal microbiota.

## Background

Malaria is still a worldwide life-threatening infection disease especially during the COVID-19 pandemic[1, 2]. In 2018, malaria caused estimated 218 million cases and 405 thousand deaths over the world. The incidence and mortality of malaria has decreased approximately 30% and 60% respectively in recent 20 years, but the disease still causes 220 cases per 1000 population at risk in WHO African region[3]. After the infection of genus *Plasmodium* and the completion of pre-erythrocytic stage development, merozoites released from hepatocytes invade red blood cells, and start the development of blood stage, which is responsible for the clinical presentation of patients, such as typical periodic fever[4].

It is well known that CD4+ T cells play critical roles in protective immunity against blood-stage *Plasmodium* infection. CD4+ T cells not only help B cells to produce parasite-specific antibody, but also enhance macrophages to kill the parasite through IFN-gamma secretion[5, 6]. However, the underlying mechanism of the activation of protective CD4+ T cell responses during *Plasmodium* infection is still largely unknown[7]. There were several studies verified that intestinal microbiota actually participates in host immune[8-10] and suggested that intestinal microbiome might be related to malaria procession. Nicolas et al[11]found that different vendors of mice with different composition of gut microbiota showed significantly different susceptibilities after infection of *Plasmodium*, while *Lactobacillus* and *Bifidobacterium* might be the important genera during the modulation. The similar findings have also been confirmed by another research in 195 Malian children[12]. With germ-free control, Mooney et al [13] analyzed fecal pellets of *Plasmodium yoelii* (*P. yoelii*) infected mice using 16s rDNA sequencing. It demonstrated that *Firmicutes/Bacteroidetes* ratio and abundance of *Proteobacteria* reduced 10 days after infection, and the unconventional state reverted to baseline by 30 days. Moreover, the colonization of *Proteobacteria Escherichia coli* and nontyphoidal *Salmonella* spp. was much easier for infected mice. For the pregnancy population, the majority of malaria patients, Catherine et al[14] showed that intestinal microbiome influences gestation outcome of mice. Bahtiyar Yilmaz et al[15] demonstrated that both *Plasmodium* spp. and human intestinal microbe *E. coli* O86:B7 could express α-gal as the anti-α-gal Abs were associated with the protection against malaria transmission in human after colonization of *E. coli* O86:B7. In addition to 16S rDNA sequencing, Stough JM and colleagues[16] combined transcriptome of whole ceca and metabolomics of contents to determine related microbiota. Moreover, Joshua ED et at[17] found that the effect of severe malaria on microbial homeostasis was greater than mild malaria using metabolomics.

According to previous studies, there was a positive relationship between intestinal microbiota and *Plasmodium* infection, however, the dynamic change of intestinal microbiome especially at species level and its relationship with gut immune response during the whole *Plasmodium* infection process are not quite clear[18-20]. Thence, the aim of our study is to investigate the potential relationship between intestinal microbiome and host immune response against *P. yoelii* infection according to the full-length 16S rDNA sequencing and intestinal transcriptome analysis.

## Results

### Parasitaemia dependent diversity and composition of mice intestinal microbiota during the whole malaria process

During the progress of *P. yoelii* infection, parasitaemia reached 10% at the 9th day, peaked around the 14th day and returned to 0 about the 20th day (Fig.1).

**Fig 1.**
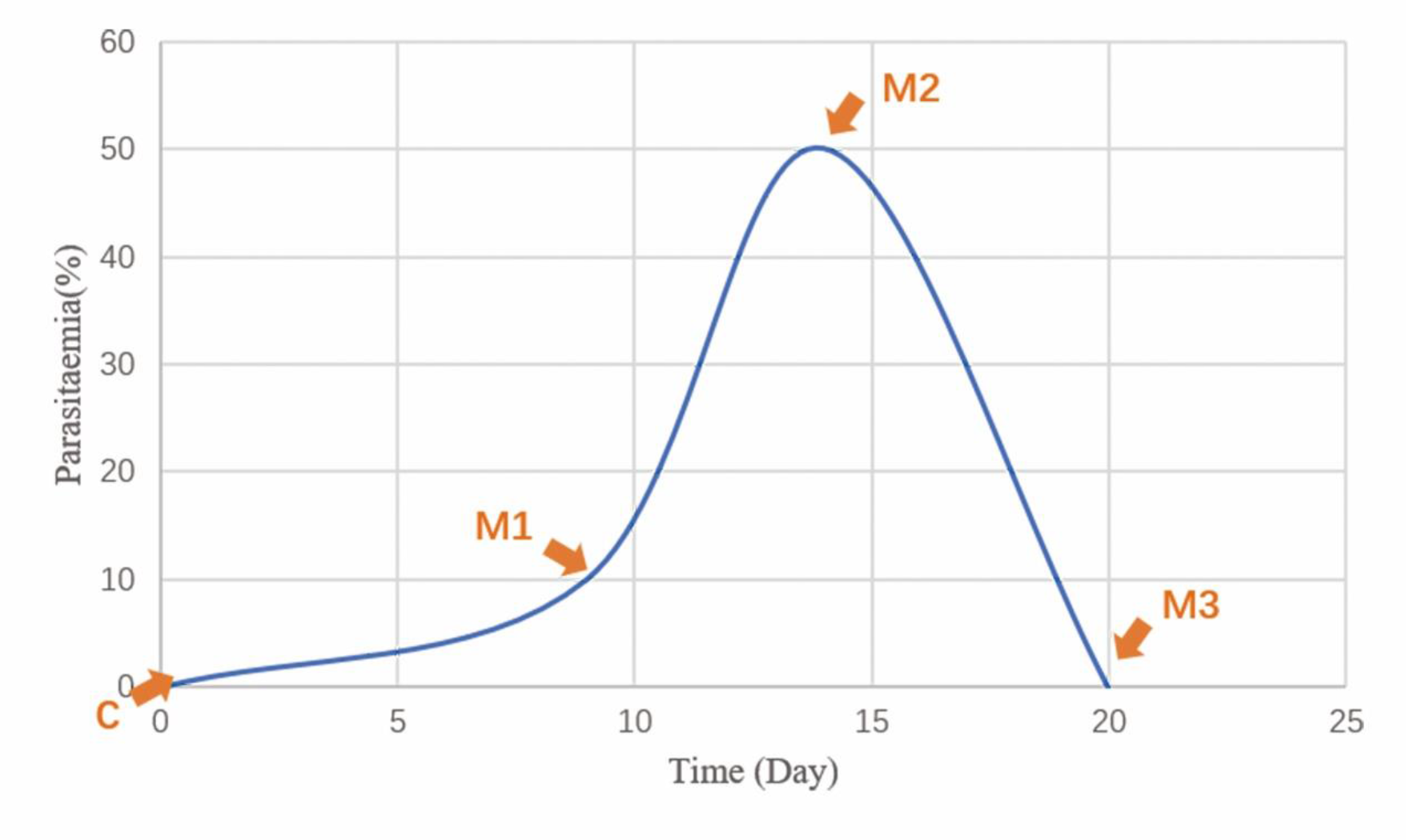
Fluctuation of Parasitaemia over infection time. Parasitaemia reached 10% at the 9th day, peaked around the 14th day and returned to 0 at about the 20th day. Arrows show time points of sample collection. Samples obtained from each time point were grouped into C, M1, M2 and M3 respectively.

We collected intestinal contents before the infection as normal control (C), 10% Parasitaemia (M1), 50% Parasitaemia (highest Parasitaemia level, M2) and 0% Parasitaemia (recovered from the infection, M3), respectively (Fig.1). Sequence data were extracted out of 15 intestinal contents samples. The amount of clean tags data was distributed between 15403-30594 after quality control and the amount of valid tags was scattered between 12653-29092 after further removal of chimera. The average length of the valid tags was 1577.14-1665 bp, and 2670-5894 OTUs were classified form each sample at 97% sequence identity.

OTU counts significantly increased over infection time (Fig.2a). However, there was not a significant decrease of OTU counts when the parasitaemia fell back (Fig.2a). There were 847 core OTUs from all of the four groups (Fig.2b). Similar with OTU counts, alpha diversity of mice intestinal microbiome increased along with parasitaemia rates (Fig.2c). Meanwhile, the community diversity and parasitaemia showed a positive correlation during the *P. yoelii* infectious phase until to the parasitaemia peak point (Fig.2d). Interestingly, the diversity recovered a little but with no statistical differences when the prarasitaema rates returned to 0% (Fig.2c) in line with the OUT counts (Fig.2a).

**Fig 2.**
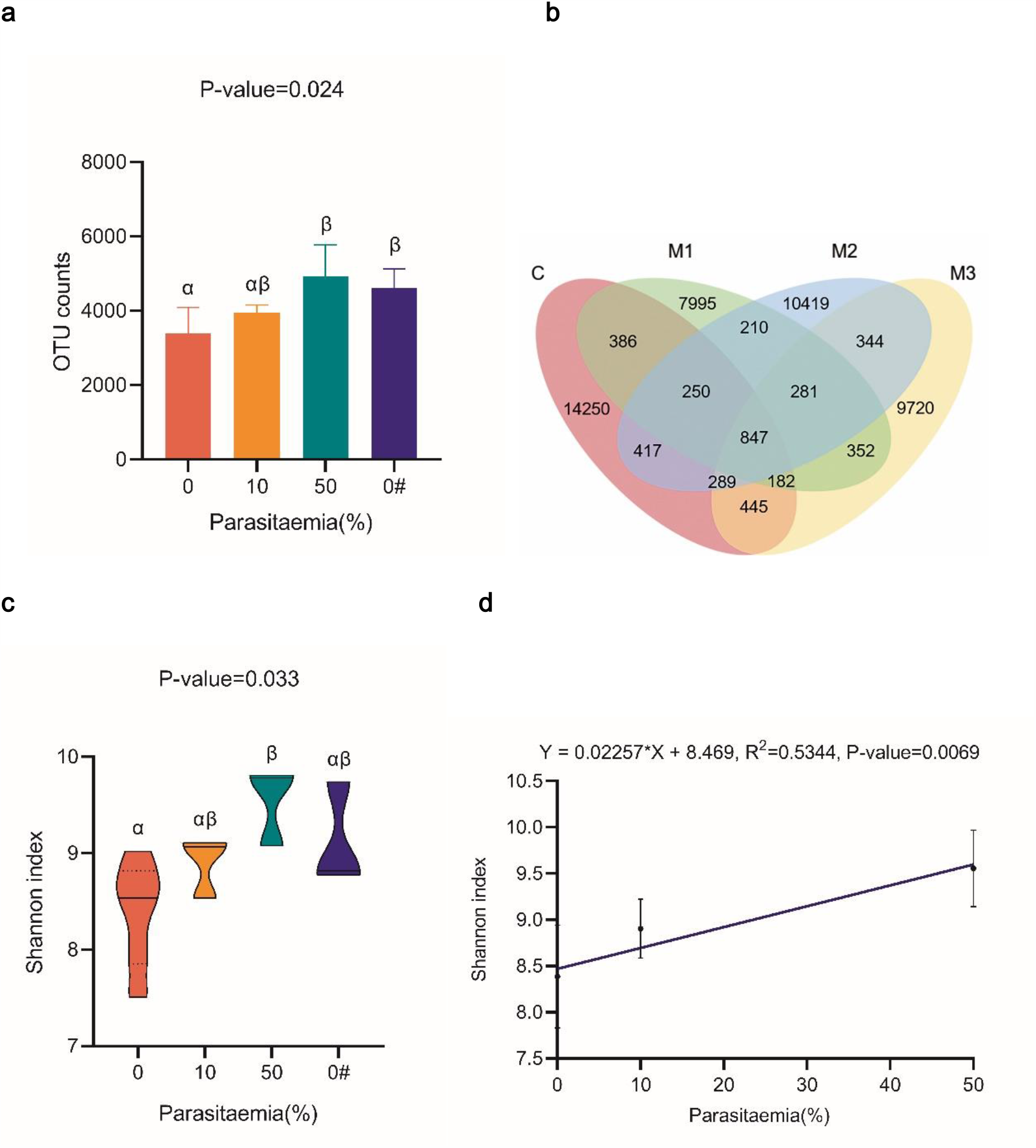
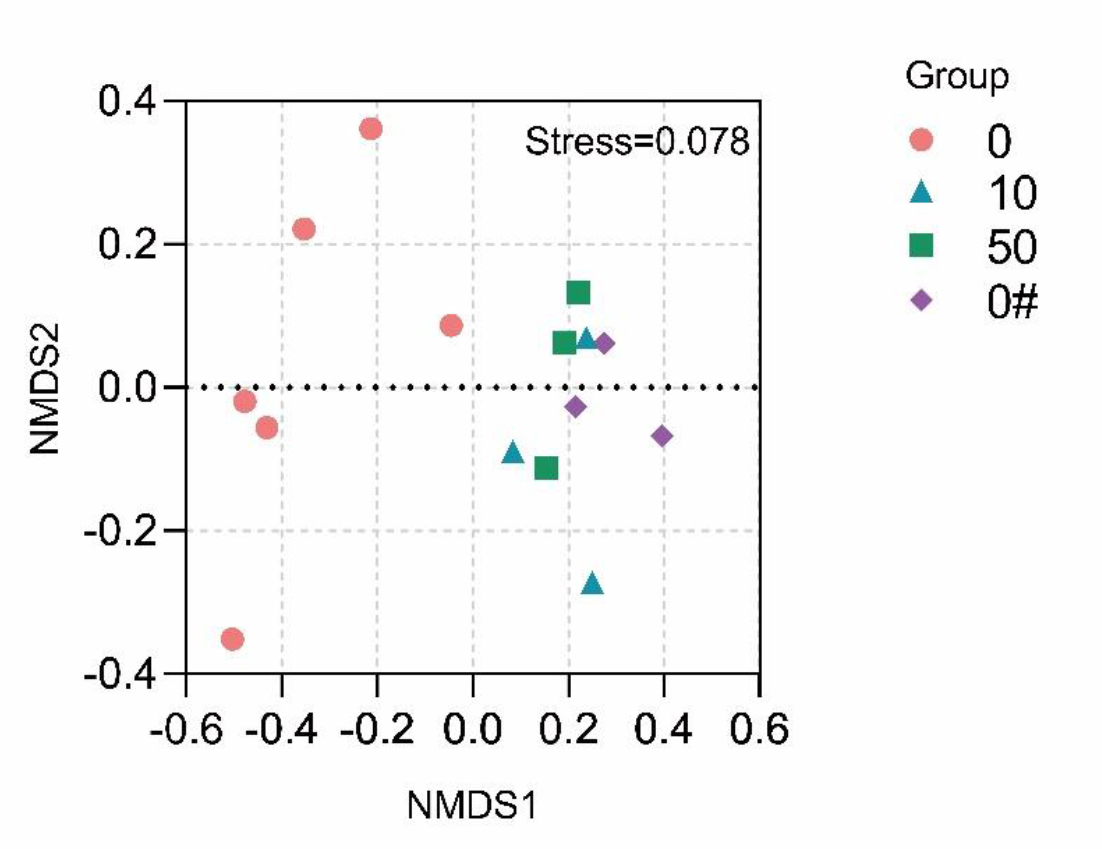
The dynamic diversity and composition of mice intestinal microbiome during the Plasmodium infection. a. OTU counts when parasitaemia was 0, 10%, 50% and returned to 0. There is significant difference (P-value < 0.05) among groups using ANOVA analysis. There is significant difference (P-value < 0.05) between groups if the column is labeled with different letters (α or β), and no significant difference (P-value > 0.05) if with the same letters (α or β). b. Venn plot of OTU taxonomy around groups. Numbers in plot: taxonomy counts of OTUs within the interval. Red: C (uninfected samples); green: M1 (10% parasitaemia); blue: M2 (50% parasitaemia); yellow: M3 (parasitaemia returned to 0). c Alpha diversity when parasitaemia was 0, 10%, 50% and returned to 0. Shannon index: alpha diversity. There is significant difference (P-value < 0.05) among groups using ANOVA analysis. There is significant difference (P-value < 0.05) between groups if the violin is labeled with different letters (α or β), and no significant difference (P-value > 0.05) if with the same letters (α or β). d Linear regression model of relationship between intestinal microbial diversity and parasitaemia. For each intestinal content sample, Shannon index was plotted against parasitaemia (x-axis). The blue line is the linear model fit. There is a significant liner slope with P-value < 0.01. e The two-dimensional NMDS plot of the microbiota samples presents similarity of OTU sequences analyzed via binary jaccard algorithm. The stress function is 0.078. Red: C (uninfected samples); blue: M1 (10% parasitaemia); green: M2 (50% parasitaemia); purple: M3 (parasitaemia returned to 0). The closer the sample clustering distance is, the more similar the samples are.

Based on binary jaccard algorithm, we analyzed microbiol composition of different groups via non-metric multidimensional scaling (NMDS). The samples were scattered in a two-dimensional plot and significantly divided into different groups in terms of parasitaemia (Fig.2e). Anosim analysis suggested that there were statistical differences of microbial composition between groups (Tab.S1). In addition, the composition of intestinal microbial community of infected mice (M1, M2 and M3 groups) was obviously different compared with uninfected mice (C group), while the structure of microbial community showed more similarities between *P. yoelii* infected groups.

### *P. yoelii* infection phase-specific Intestinal microbiota characteristics at family, genus and species levels

At the family level, only the abundance of *Rikenellaceae*, Family XIII, *Peptococcaceae* and *Bacteroidales* BS11 gut group showed differences (Tab.S2). It was notable that the abundance of *Rikenellaceae* gradually reduced after *P. yoelii* infection, while *Bacteroidales* BS11 gut group could only be examined in acute infection phase (∼10% parasitaemia) (Fig.3a). The abundance of *Peptococcaceae* reached to the maximum along with the rising of parasitaemia and the minimal abundance of Family XIII was in the malaria recovering phase (Fig.3a).

**Fig 3.**
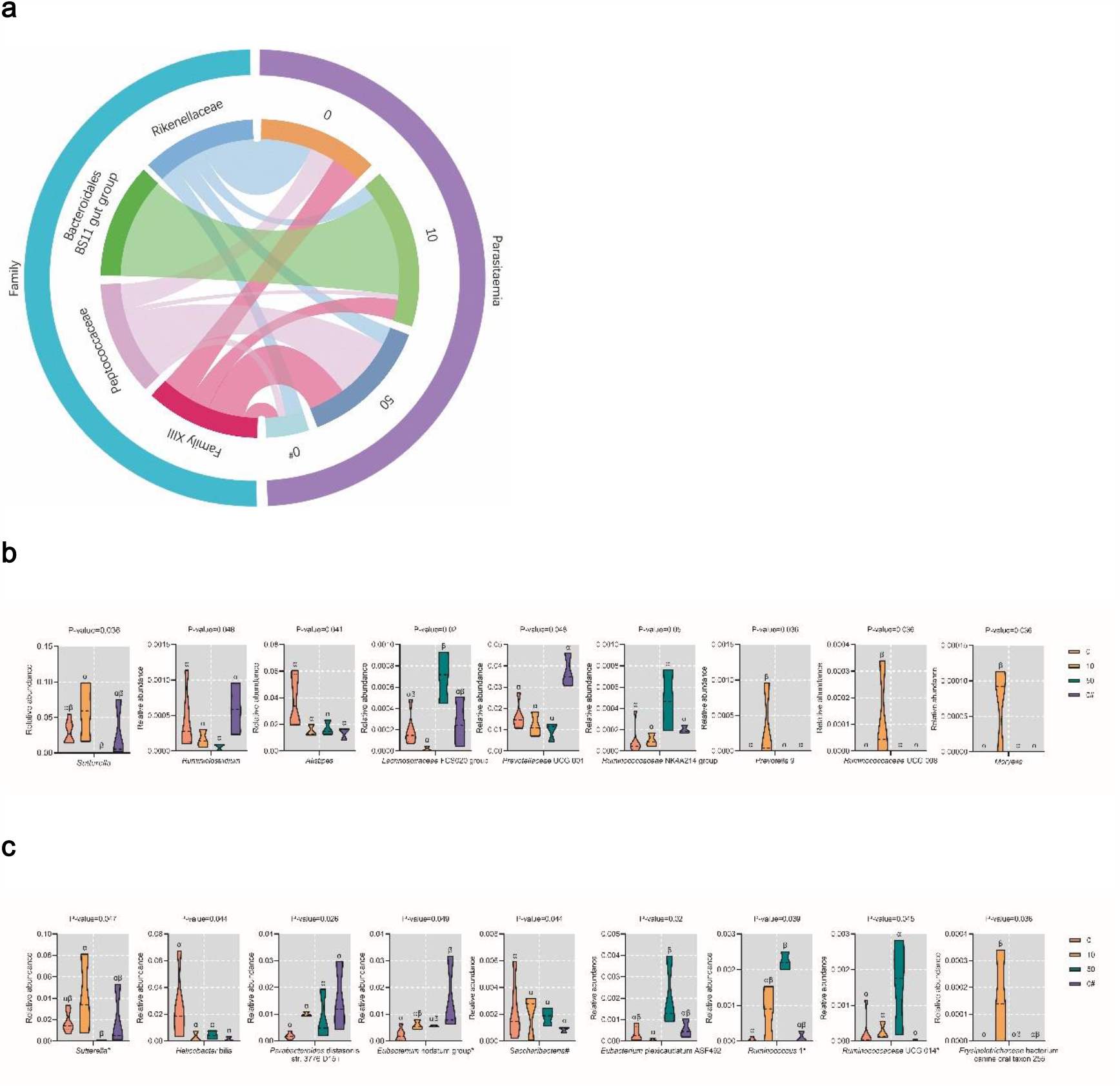
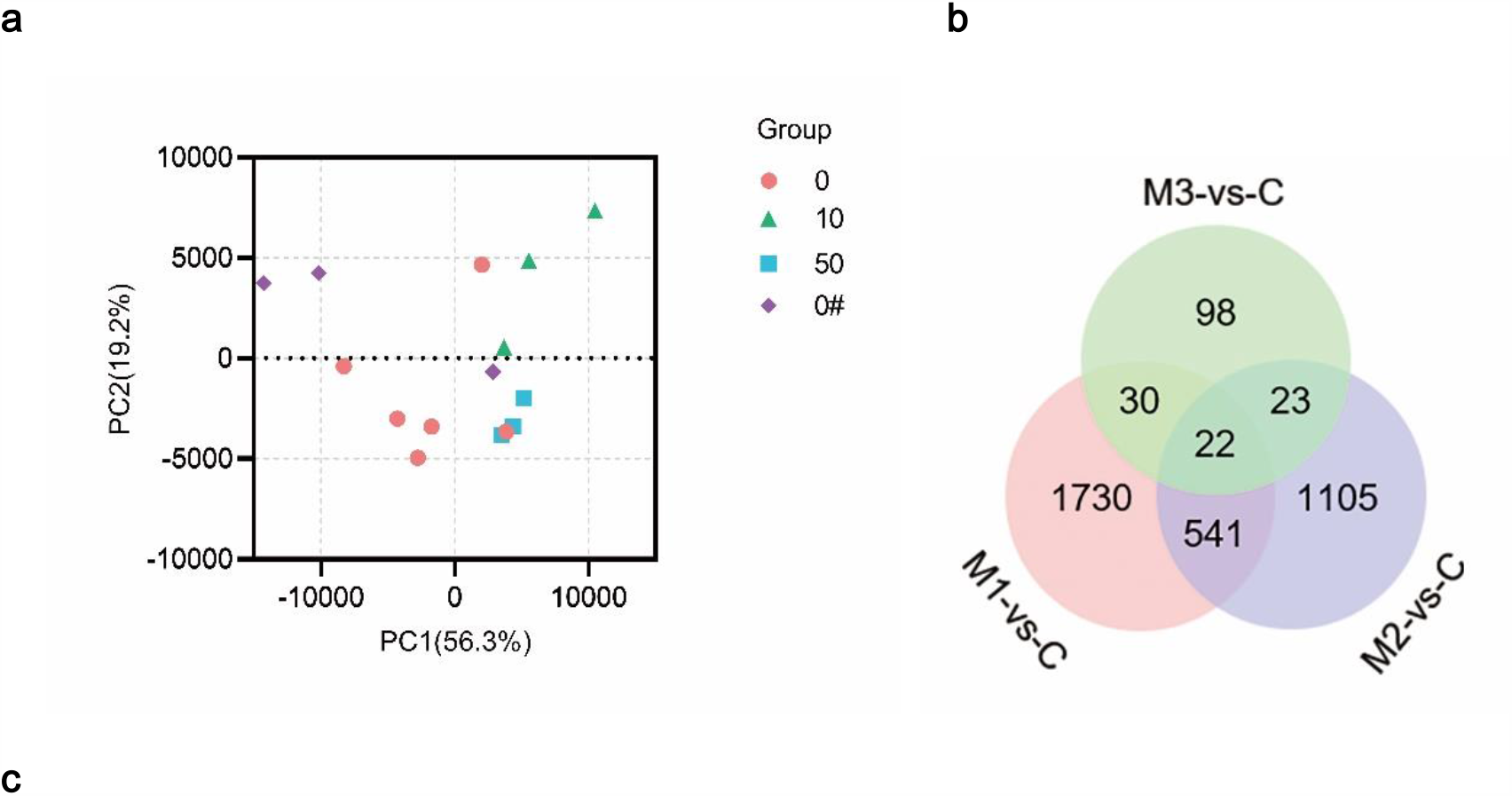
Microbial structure changes during the malaria procession. a Percentage of microbiota abundance in family level. The left semicircle shows microbiota with different abundance and the right semicircle presents malaria procession by parasitaemia. Families and parasitaemia are identified via colors. The width of colored stripes reflects percentage of abundance in certain malaria phase. b Relative abundance of differential microbiota in genus level. Different violins represent different groups. c Relative abundance of differential microbiota in species level. Different violins represent different groups. Red: uninfected; yellow: 10% parasitaemia; green: 50% parasitaemia; blue: parasitaemia returned to 0. There is significant difference (P-value < 0.05) among groups if the data is labeled with different letters (α or β), and no significant difference (P-value > 0.05) if with the same letters (α or β) in the same graph. * Unknown species of corresponding genus. # Unknown species of corresponding phylum.

At the genus level, the abundance of *Sutterella, Ruminiclostridium* and *Prevotellaceae* UCG 001 dropped to the minimum while the abundance of *Lachnospiraceae* FCS020 group and *Ruminococcaceae* NK4A214 group reached to the maximum along with the rising of parasitaemia (Fig.3b). Meanwhile, the genera *Prevotella* 9, *Ruminococcaceae* UCG 008 and *Moryella* can only be detected when the parasitaemia reached to 10% (Fig.3b). Besides, the abundance of *Alistipes* reduced into a stable line after the infection (Fig.3b).

As in Fig.3c, at the species level, *Erysipelotrichaceae* bacterium canine oral taxon 255 could only be classified in acute *P. yoelii* infection (Fig.3c). Abundance of *Eubacterium* plexicaudatum ASF492, *Ruminococcus* 1* and *Ruminococcaceae* UCG 014* peaked matched with 50% parasitaemia (Fig.3c). However, *Sutterella*\* presented an opposite trend with the minimum abundance in peak of parasitaemia (Fig.3c). Along with the whole disease procession, the abundance of *Parabacteroides* distasonis str. 3776 D15 i and *Eubacterium* nodatum group* kept rising while the abundance of *Saccharibacteria*^*#*^ gradually reduced (Fig.3c). Different with above trends, *Helicobacter* bilis could only be detected in a lower abundance after *P. yoelii* infection (Fig.3c).

### The shift of intestinal transcriptome in the response to *P. yoelii* infection

In order to explore how the intestine responded to *P. yoelii* infection, RNA was extracted from mice colorectal and analyzed by using transcriptome sequencing. A total of 136.7G of clean data was obtained and the clean base of each sample was distributed from 6.78G to 8.28G after quality preprocessing. In addition, the Q30 base was scattered from 92.28% to 94.37% while the average GC content was 48.95%. Then, clean reads were performed sequence alignment with the reference genome. The comparison rates were 78.53%-97.50%. The correlation coefficient from the sequencing samples based on gene expression revealed that samples were in good reliability and rationality (Fig.S1). According to the Principal component analysis (PCA analysis), malaria procession significantly affected the mice intestinal gene expression (Fig.4a).

**Fig 4.**
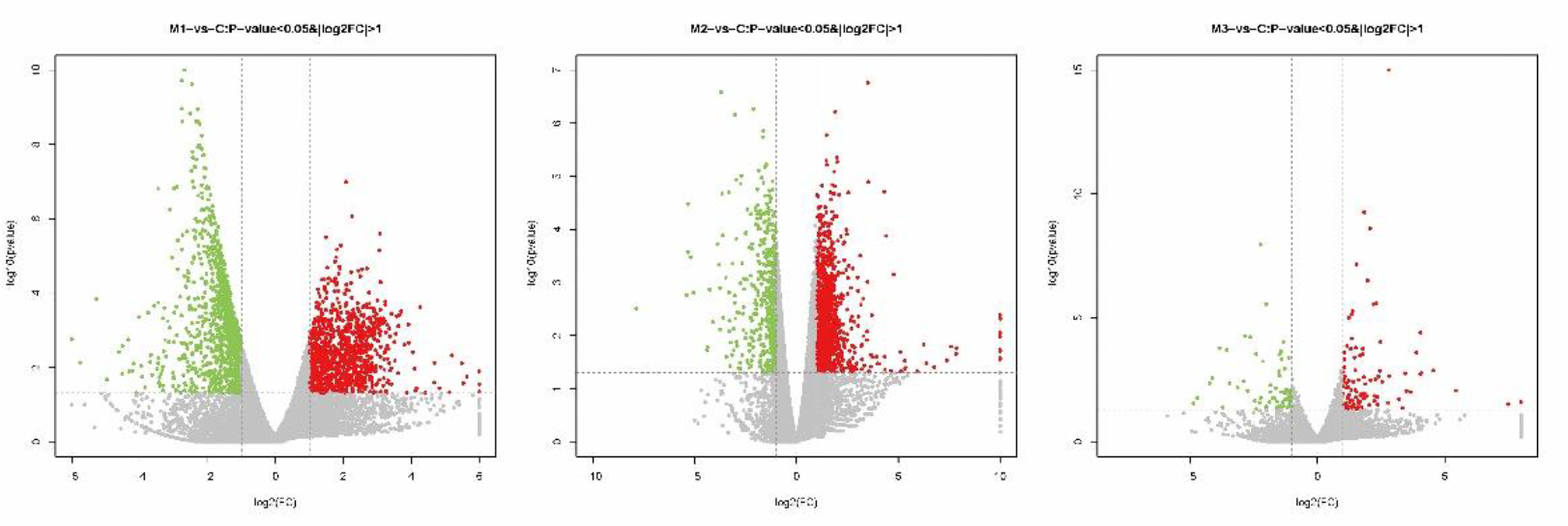
Intestinal genes expressed differently along with the malaria procession. a Principal component analysis (PCA analysis) of mice intestinal gene expression among groups with malaria procession. Red: C (uninfected samples); green: M1 (10% parasitaemia); blue: M2 (50% parasitaemia); purple: M3 (parasitaemia returned to 0). The closer the sample clustering distance is, the more similar the samples are. b Venn plot of differently expressed genes along with malaria procession. Numbers in plot: counts of differently expressed genes. Red: M1 compared with C group; blue: M2 compared with C group; green: M3 compared with C group. c Volcano plots of differently expressed genes filtering by P-value < 0.05 and |log2FC| >1. X axis: log2 fold change in gene expression; Y axis: -log10 P value. Red: up-regulated genes; green: down-regulated genes; gray: genes with non-significantly different expression.

There were 22 core genes representing significant difference along with the malaria process (Fig.4b).The number of the differently expressed genes at 10 % and 50% parasitaemia was significantly increased comparing with uninfected samples, while was reduced when the parasitaemia recovered to 0% (Fig.4c). There were 42.36% and 72.32% of those genes up-regulated when parasitaemia was at 10% and 50% respectively (Fig.S2). The similarity between M3 group (recovered group) and C group (uninfected group) was extremely higher than that from M1 (10% parasitaemia) and M2 (50% parasitaemia) groups (Fig.S3), indicating that the intestinal response was highly corelated to the infectious states.

### The dynamic change of host immunity related genes highly correlated with the parasitaemia states

There were 120, 49 and 58 pathways with statistically significance when parasitaemia reached 10% (M1), 50% (M2) and returned to 0 (M3) respectively. Interestingly, the metabolism-related pathways were mainly down-regulated, while the up-regulated pathways were mostly related to the host immunity, such as Th1 and Th2 cell differentiation, Th17 cell differentiation, T cell receptor signaling pathway, B cell receptor signaling pathway and intestinal immune network for IgA production (Fig.5a and Fig.S4). It suggested that these immune pathways participated in different parasitaemia states. Among the 63 up-regulated genes from the above enriched pathways, the expression of 15 core genes (Bcl6, Cd40lg, Gata3, Icos, Il10, Il12a, Il12b, Il21, Il4, Maf, Rorc, Stat4, Tbx21, Tgfb1 and Tnfrsf4) represented a state-specific characteristic (Tab. S3, Tab.1 and Fig.5b).

**Fig 5.**
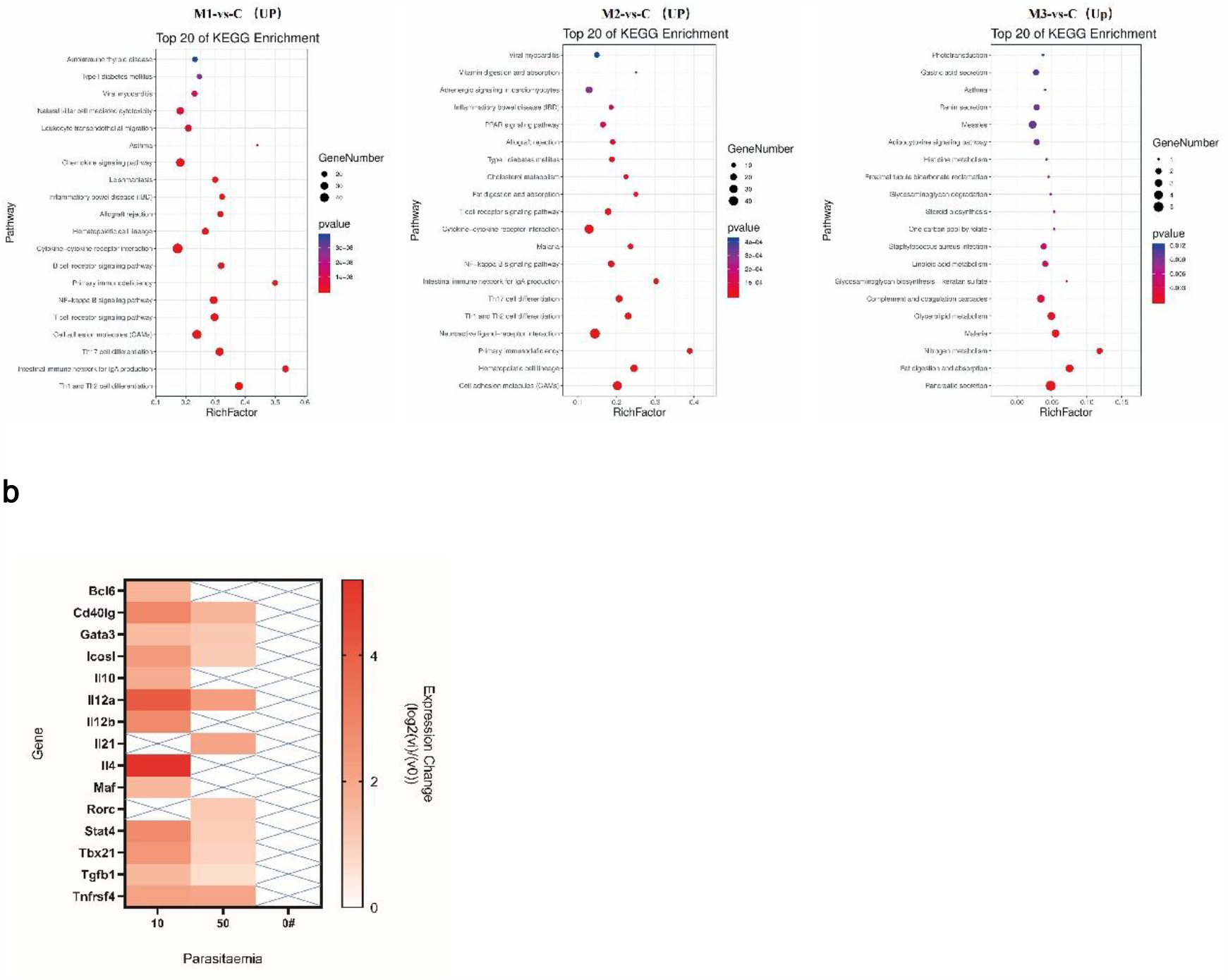
Up-regulated genes are mainly enriched in immune-related pathways. a Top 20 pathways of up-regulated genes by KEGG enrichment. The bubble chart indicates gene pathways enriched via KEGG analysis. The X axis shows rich factor while the left vertical axis lists different gene pathways. The larger the bubble is, the more the number of differential protein coding genes contained in the entry. The P-value legend reveals the degree of enrichment significance. The bubble color changes comply red to blue while P-value increasing. b Change of gene expression compared with uninfected samples. X axis: parasitaemia. Left Y axis: gene id. Legend: expression change (log2(vi)/(v0)). Red: up-regulated gene with P-value < 0.05; ×: gene with no significant difference of expression (P-value > 0.05).

**Table 1.** Core genes of up-regulated immune pathways

| Core gene | Gene function |
| --- | --- |
| Bcl6 | Coding BCL6, the master transcription factor of Tfh differentiation |
| Cd40lg | Coding CD40L, the activation cytokine of B2 cell differentiation |
| Gata3 | Coding Gata3, the master transcription factor of Th2 cell differentiation |
| Icosl | Coding ICOS, the promoting factor of Tfh and the signal transfer between Tfh and B cell |
| Il10 | Coding IL-10, the activation cytokine of B1 cell and plasma cell differentiation |
| Il12a | Coding IL-12, the activation cytokine of Th1 cell differentiation and the signal transfer between Tfh and B cell |
| Il12b | Coding IL-12, the activation cytokine of Th1 cell differentiation and the signal transfer between Tfh and B cell |
| Il21 | Coding IL-21, the activation cytokine of Th17 cell differentiation and the main secretory factor of Tfh |
| Il4 | Coding IL-4, the activation cytokine of Th2 cell and B1/B2 cell differentiation |
| Maf | Coding c-Maf, the master upstream activator of IL-4 production |
| Rorc | Coding RORγt, the master transcription factor of Th17 cell differentiation |
| Stat4 | Coding STAT4, the master upstream activator of T-bet and IFN-γ production |
| Tbx21 | Coding T-bet, the master transcription factor of Th1 cell differentiation |
| Tgfb1 | Coding TGF-β, the secretory factor of Th17 cell and the activation cytokine of B1/B2 cell differentiation |
| Tnfrsf4 | Coding OX-40, the promoting factor of Tfh and the signal transfer between Tfh and B cell |

#### 1. Th1 and Th2 cell

Th1 and Th2 cell is a crucial differentiation direction of CD4+ T cell in antimalarial procession[21]. Comparing with uninfected group, Il12a, Il12b, Stat4 and Tbx21 were significantly up-regulated until parasitaemia reached to the peak (Tab.1, Tab.S3 and Fig.5b). However, the expression of the above genes showed a downward trend along with the malaria process (Fig.5b) indicating that IFN-γ+ Th1 cell, the early immune responder, was gradually depleted over time. Core genes, including Il4, Maf and Gata3, were mainly expressed at 10% parasitaemia (Tab.1, Tab.S3 and Fig.5b) suggesting that Th2 might only appear in early host immune response after *P. yoelii* infection.

#### 2. Th17 cell

Th17 cell was only activated at 50 parasitaemia according to the up-regulation of Il21 and Rorc genes (Tab.1, Tab.S3 and Fig.5b) indicating that Th17 cell was an essential immune executor with TGF-β production.

#### 3. Follicular helper T cell (Tfh)

Overexpression of BCL6, the master transcription factor of Tfh differentiation, only appeared in early immune response (Tab.1, Tab.S3 and Fig.5b). However, there was a consistent up-regulation of promoting factors including IL-12, IL-21, ICOS and OX-40 (Tab.1, Tab.S3 and Fig.5b) suggesting the early occurrence of related antibody and its sustained effect before the total clearance of *P. yoelii*. Moreover, the activities of all of the above CD4+ T cells recovered to the uninfected levels when the parasitaemia returned to 0.

#### 4. B cell

Activators of Peyer’s patch from B1/B2 cells, including CD40L, IL-4, IL-10 and TGF-β, were consecutively up-regulated along with the parasitaemia (Tab.1, Tab.S3 and Fig.5b). Cooperating with phase specific activation of Th1/Th2 cells, Th17 and Tfh cells, B cells could be active in the whole host immune response to *P. yoelii* infection. After the recognition of signals from multi-differentiated CD4+ T cells and the plasma cell-directed differentiation, B cells can produce malaria-specific antibody (such as intestinal IgA) against *P. yoelii* infection. Then after the *P. yoelii* clearance, these genes returned to the uninfected levels, indicating that the active B cells was also depleted until the parasitaemia recovered to a safe titer.

The host immune cells represented a parasitaemia phase-specific characteristics against *P. yoelii* infection (Fig.6). In the early immune response, naive CD4+ T cells were activated to differentiate into IFN-γ+ Th1, IL-4+ Th2 cells and IL-21+ Tfh cells after the antigen presentation. Meanwhile, B cells started to interact with the helper T cells and secret effective antibody (intestinal IgA etc.). Along with the rising of parasitaemia, assisters of B cells were gradually replaced from Th1/Th2 and Tfh cells to Th1, TGF-β+ Th17 and Tfh cells. The process was accompanied with the reduction of the activity from Th1/Th2 cells and the continuous activation of B cells. Moreover, all the above activated immune cells were returned to the uninfected levels after the accomplishment of the clearance of *P. yoelii*.

**Fig 6.**
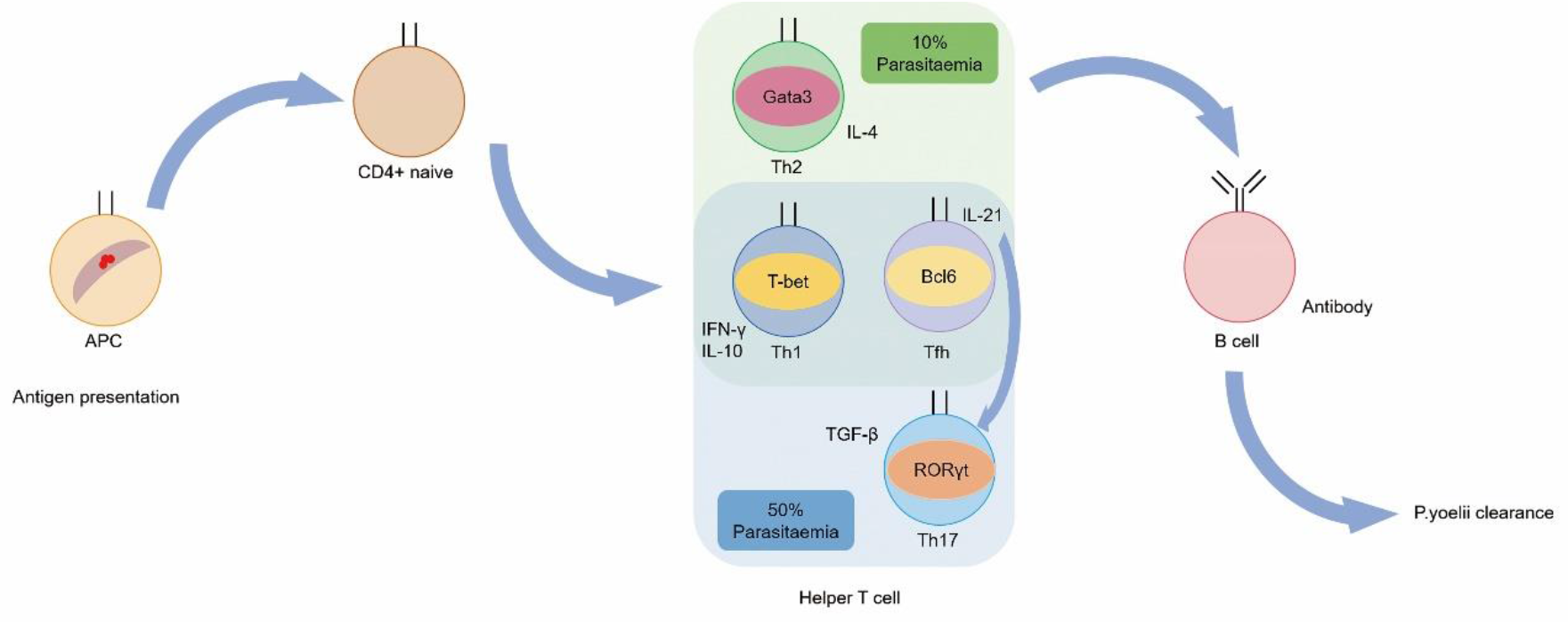
Graphic of host immune response against *P. yoelii* infection.

### Intestinal microbiota was correlated with the host immune response against *P. yoelii* infection

We then analyzed the relationship between intestinal microbiota at family, genus and species levels and host immunity by calculating person index (Fig.7). At family level, *Bacteroidales* BS11 gut group was only detected at 10% parasitaemia (Fig.3a) and it was in a high-correlation with the expression of Cd40lg, Icos, Il10, Il12a, Il12b and Stat4 (Fig.7) indicated that *Bacteroidales* BS11 gut group was related to early host immune response by activating Th1 cells and signal transferring with B cells. The genera *Prevotella* 9, *Ruminococcaceae* UCG 008 and *Moryella* presented similar correlation at genus level (Fig.7), while *Moryella* and Bc16 also showed strong correlation, indicated that IL-21+ Tfh cells also participated in the early host immune response to parasitaemia. Rorc expression was in a high-correlation with the family *Peptococcaceae* and genus *Lachnospiraceae* FCS020 group whose abundance were significant higher in 50% parasitaemia (Fig.3b, 3c and Fig.7), indicated that *Peptococcaceae* and *Lachnospiraceae* FCS020 group were related with Th17 cells in later host immune response against *P. yoelii* infection.

**Fig 7.**
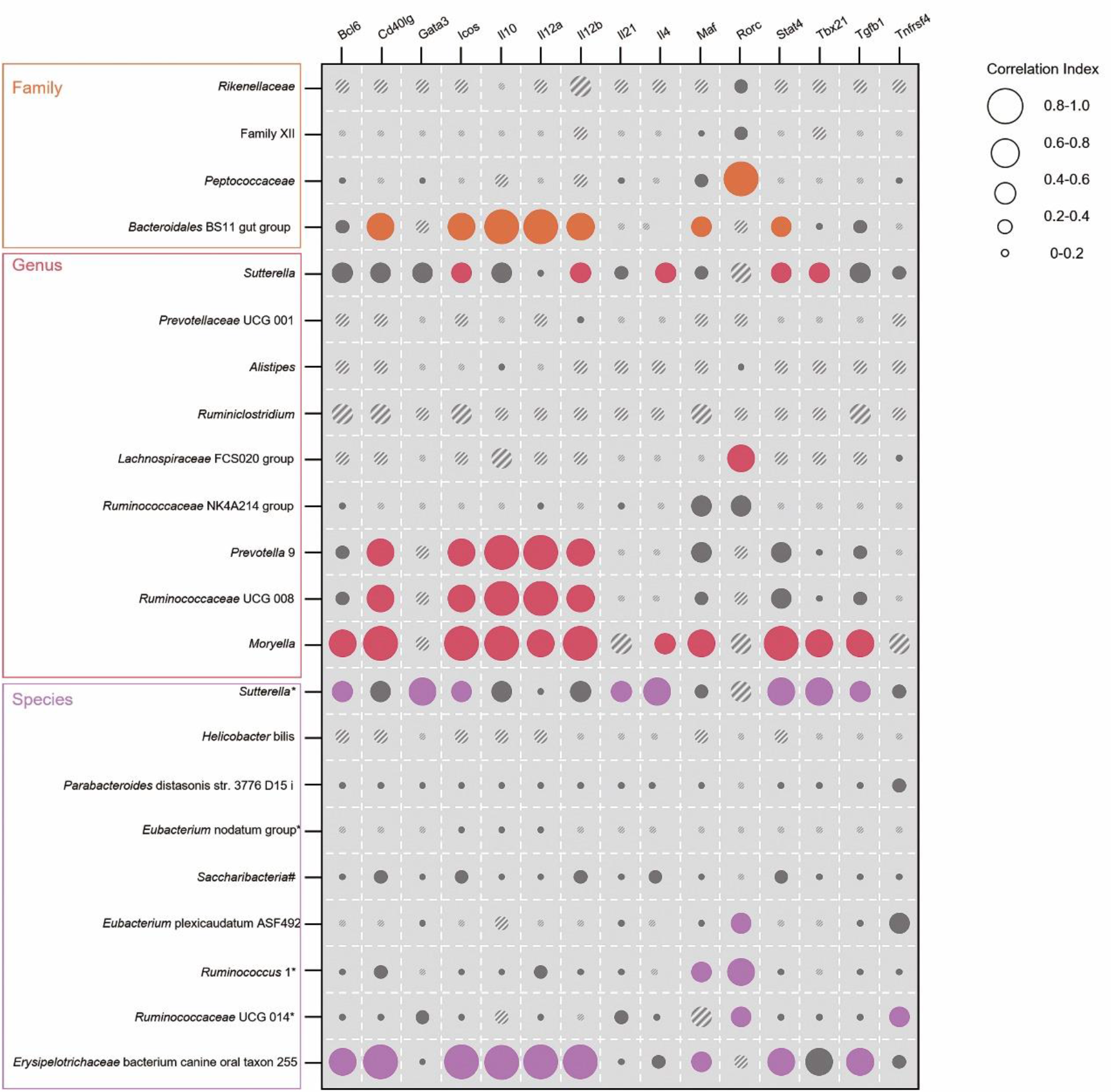
Correlation between microbiota from Fig.3b,3c and genes from Fig.5a. Only the microbiota with statistical correlation index (P-value < 0.05) are represented with colored circles (orange, red and purple) while the circles were in grey when p > 0.05. Solid circles present positive correlation while them with slashed strips relate to negative correlation. Circle size indicates the correlation index in samples, and the bigger circle means the higher index.

The genus *Sutterella*\* was high-correlated with Stat4, Tbx21, Gata3 and Il4, genes coding master transcription factors of IFN-γ+ Th1 and IL-4+ Th2 cells differentiation (Fig.7). The abundance of *Sutterella*\* significantly reduced from 10% to 50% parasitaemia (Fig.3c) suggested that the early activation of Th1/Th2 cells and the reduce of IFN-γ and IL-4 along with the development of parasitaemia were related to the abundance change of *Sutterella\**.

Both the genus *Moryella* and the species *Erysipelotrichaceae* bacterium canine oral taxon 255 were in a high correlation with Tfh and B cells activation in early host immune response (Fig.7). Another several species, including *Ruminococcus* 1*, *Ruminococcus* UGG 014* and *Eubacterium* plexicaudatum ASF492, were highly correlated with Rorc expression (Fig.7) and the abundance of these three species was dominant when the parasitaemia reached to the peak (Fig.3c).

## Discussion

Our research for the first time revealed the dynamic interactions between the intestinal microbiota at species level by using full-length 16S rDNA sequencing and the full process of *P. yoelii* infection. Along with the rising of parasitaemia, Shannon index of alpha diversity of intestinal microbiota represented a positive liner correlation. At different phase of malaria infection, intestinal microbial communities were significantly changed, specifically, in family level, the abundance of *Rikenellaceae* significantly reduced along with the infection of *P. yoelii* while *Prevotella* 9, could only be identified in genus level in the same state. At about 14 days after infection, parasitaemia reached to the peak and the abundance of the genus *Lachnospiraceae* FCS020 group increased to the maximal level. In contrast, the abundance of *Sutterella* displayed oppositely along with the parasitaemia change.

The host response was also correlated to *P. yoelii* infection according to the intestinal transcriptome. The counts of differently expressed genes were higher in pathological stage than uninfected and recovered stages. Especially, genes related to host immune response were significantly up-regulated at 10% and 50% parasitaemia according to the KEGG enrichment analysis. It suggested that IFN-ϒ+ Th1 cells, IL-4+ Th2 cells and IL-21+ Tfh cells were early activated in helping B cells. TGF-β+ Th17 cells gradually replaced Th2 cells to cooperate with other CD4+ T cells in the later host immune response. Meanwhile, B cells could maintain activity during the whole process. The down-regulated pathways during the infection were mainly associated with the metabolism (Fig.S4), which might be caused by the shifted microbiome and the activation of host immune[22].

The intestinal microbiota was also highly correlated with the activation of certain immune cells. Especially several microbes at species level, such as *Erysipelotrichaceae bacterium* canine oral taxon 255 and *Eubacterium plexicaudatum* ASF492, were in a high correlation with activating Th1, Th2 and Th17 cells. Our results might provide a novel biomarker for malaria diagnosis and therapy by analyzing intestinal microbiome.

Previous studies of the relationship between intestinal microbiome and malaria have found several related microbes at different levels. Different from our study, Mooney et al[13] operated germ-free control before *P. yoelii* infection. They found that the abundance of *Firmicutes* reduced while the abundance of *Bacteroidetes* significantly increased at 10 days after infection, while the shifted intestinal microbial community could recover to uninfected stage by 30 days. Some others found that the abundance or colonization of *Actinobacteria* and *Proteobacteria* were related to severity of malaria at phylum level[11, 12, 15, 23]. Although the intestinal microbial community was shifted after *P. yoelii* infection, there was no any difference in phylum level during malaria process in our study. However, we identified different microbes at genus and species levels along with the *P. yoelii* infection process.

We then invested the host response to *P. yoelii* infection and its relationship with intestinal microbiota. Previously, Stough JM et al[16] combined transcriptome and metabolomics to investigate microbial gene expression and their functions during parasite infection. In our study, we combined intestinal microbiome and transcriptome to determine correlation between the microbial abundance change and immune gene expression during the whole *P. yoelii* infection process, and we identified the different microbiota at species level and their correlation with host immune response. Villarino N.F. et al[11] applied fecal transplant to verify the relationship between intestinal microbiome and malaria. Based on our results, specific microbes at species level, such as *Erysipelotrichaceae* bacterium canine oral taxon 255, *Ruminococcus* 1*, *Ruminococcus* UGG 014* and *Eubacterium* plexicaudatum ASF492, can be transplant to explore more accurate investigation of host response or therapeutic targets.

*P. yoelii* can be eliminated by host immune [24]. CD4+ T cells have been proved to play a central role in immune protection against *P. yoelii* infection [21]. Th1 cells are considered as a crucial part of acute control of parasite infection through the production of IFN-γ [25-27]. Our results proved that Th1 cells played an important role during acute immune response with up-regulated T-bet. However, Oakley et al[28] found that malaria could be controlled even in T-bet deficient mice. In some infection models, Th2 cells were in charge of activating B cells in chronic infection phase[26, 27]. Differently, we found that the activity of Th2 cells was depleted when parasitaemia reached to the peak and Th17 cells replaced them to assist B cells in protecting host from infected stage. There are some other CD4+ T cell sets also participating in host immune against *P. yoelii* infection. The positive function in controlling malaria of Th17 cells is still controversial[21, 29, 30]. In our results, Th17 cells were activated by IL-21 when the parasitaemia reached to the peak. IL-21 is an essential production of Tfh cells. However, there were few data on Tfh cells in *P. yoelii* infection[31]. It has been demonstrated that IL-21 response is enhanced in blood of immune adults living in endemic areas of *P. falciparum* transmission[32, 33]. Tfh cells can also modulate IgA secretion cooperating with Th17 cells[34]. Our results showed that IL-21-producing Tfh cells were activated in earlier host immune response. Similar with chronic immune function of Th17 cells in extracellular pathogens, such as HIV[35], we supposed that Th17 cells took part in the later host immune response against high titer parasite infection. According to our results, the function of immune cells during malaria process is dynamic and interactional. Meanwhile, it’s worth to noticed that genes coding major histocompatibility complex (MHC), surface molecule on antigen-presenting cells (APC), also kept at a high level of expression (Tab.S3) although all the immune cells we found returned to the uninfected level after the *P. yoelii* clearance. It’s benefits to the host might cause a faster immune response and explain the resistance when exposed to the next infection[36].

We also found a high correlation between certain intestinal microbiota and master transcription factors of host immunity during *P. yoelii* infection. Currently, we still cannot identify their causality[37]. The intestinal microbiota can produce serval metabolic repertoire as signal molecules, such as short-chain fatty acids (SCFAs), or increase the host product, such as bile acid, to regulate the phenotype of anti-inflammatory cells [38-40]. Meanwhile, host immunity can also regulate the characteristic of microbiota through their interaction such as the host immunity-microbiota interaction in inflammatory bowel disease (IBD) [40]. We found that the intestinal microbiota was related to host immune response against *P. yoelii* infection according to our results and we will investigate which is the first causality whether the shift of intestinal microbiota combined *P. yoelii* infection activates the host response or the host response caused by *P. yoelii* infection drives the change of microbiota in further researches. Additionally, some studies were trying to explain the negative correlation between malaria prevalence and mortality of certain cancers [41, 42]. According to our results, the up-regulated systemic immune by intestinal microbiome might affect cancer clinical outcomes [43–45].

In summary, Our findings, that the shift of intestinal microbiota was highly related to immune cells to protect host from *P. yoelii* infection, highlighted the critical roles of intestinal microbiota in *P. yoelii* infection and host immune response, which provided new information to understanding the pathogenesis of *Plasmodium* and treatment.

## Materials and Methods

### Mice and Infections

BALB/c mice at age of 6-8week were provided from the Experimental Animal Center of the Third Military Medical University. Mice were infected by injection intraperitoneally with 100μl inoculum containing 10^5^ *P. yoelii* 17XNL parasitized red blood cells from the donor mice. The parasitemia was detected by Gemisa staining and recorded every 2 days.

### Collection of colorectal and its contents

Before the infection, colorectal and its contents were collected from C group mice as control samples. The colorectal and its contents were collected from the remaining mice when parasitemia reached 10%, 50% and returned to 0, and labelled as M1, M2 and M3 group, respectively.

### Gut Microbiota analysis

Gut microbiota analysis was accomplished by Shanghai OE Biotech Company. The colorectal contents were flash-frozen and stored in liquid nitrogen. Microbial DNA was isolated by using the ZymoBIOMICS DNA Miniprep Kit (Zymo Research, Irvine, California, USA) in a biological flow cabinet. DNA sequencing was completed using SMRT provided by PacBio Sequel. Vsearch software package was used to get phylogenetic tree file and OTU classification table compared with Silva database (v123)[46]. The Shannon-Wiener index was calculated to describe alpha diversity of microbial community. Based on binary jaccard algorithm, Nonmetric Multidimensional Scaling (NMDS) was used to analysis beta diversity and Anoism analysis was applied to calculated differences. Microbial abundance data with non-normal distribution was performed Kruskal Wallis test.

### Intestinal RNA transcriptome analysis

Intestinal RNA transcriptome analysis was accomplished by Shanghai OE Biotech Company. The colorectal of mice was flash-frozen in liquid nitrogen and chose as the analysis sample. Total RNA was extracted using the mirVana miRNA Isolation Kit (Ambion) following the manufacturer’s protocol. RNA integrity was evaluated using the Agilent 2100 Bioanalyzer (Agilent Technologies, Santa Clara, CA, USA). The samples with RNA Integrity Number (RIN) ≥ 7 were subjected to the subsequent analysis. The libraries were constructed using TruSeq Stranded mRNA LTSample Prep Kit (Illumina, San Diego, CA, USA) according to the manufacturer’s instructions. Then these libraries were sequenced on the Illumina sequencing platform (HiSeqTM 2500 or Illumina HiSeq X Ten) and 125bp/150bp paired-end reads were generated. Trimmomatic[47] was used to perform quality control. Then the clean reads were mapped to reference genome using hisat[48]. FPKM[49] value of each gene was calculated using cufflinks[50] and the read counts of each gene were obtained by htseq count[51]. DEGs were identified using the DESeq[52] (2012) R package functions estimate Size Factors and nbinom Test. Hierarchical cluster analysis of DEGs was performed to explore genes expression pattern. KEGG[53] pathway enrichment analysis of DEGs were respectively performed using R based on the hypergeometric distribution. STEM software package was used to describe gene expression fluctuation during malaria procession.

### Statistical Analysis

Descriptive and comparative statistical analyses of data, except the gut microbiota and intestinal transcriptome data, were done using IBM SPSS Statistics 23 and GraphPad Software (Prism, version 8).

## Supplementary information

**Additional file1: Table S1**. Anosim analysis of NMDS. **Table S2**. Kruskal-Wallis Test of intestinal microbial relative abundance on family level. **Figure S1**. Inter-sample correlation. The abscissa represents the sample name, and the ordinate represents the corresponding sample name. Blue: samples with high correlation coefficient; white: low correlation coefficient among samples. **Table S3**. Expression of genes related to host immunity with statistical differences. **Figure S2**. Venn plot of differently expressed genes along with malaria procession. Numbers in plot: counts of differently expressed genes. Red: M1 compared with C group; blue: M2 compared with C group; green: M3 compared with C group. **Figure S3**. Similarities and differences between groups. Heatmap shows clustering analysis on unsupervised level (P-value < 0.05 and |log2FC| > 1) of differently expressed genes. X-axis: samples. Legend: expression change (log2(vi)/(v0)) (Red: up-regulated gene; blue: down-regulated gene). (a) M1 compared with C. (b) M2 compared with C. (c) M3 compared with C. **Figure S4**. Top 20 pathways of down-regulated genes by KEGG enrichment. The bubble chart indicates gene pathways enriched via KEGG analysis. The X axis shows rich factor while the left vertical axis lists different gene pathways. The larger the bubble is, the more the number of differential protein coding genes contained in the entry. The P-value legend reveals the degree of enrichment significance. The bubble color changes comply red to blue while P-value increasing.

## Authors’ contributions

RB, ZXD, CL, and XWY developed the original concepts. ZYW and CXY collected samples. ZYW, CL, LBY, LTP and LJY performed statistical analyses. ZYW, CL, CXY LBY, YXC and LTP wrote the paper. ZXD, XWY, and RB edited the manuscript. All authors were given the opportunity to review the results and comment the manuscript. All authors read and approved the final manuscript.

## Funding

This work was supported by National Natural Science Foundation of China grant: 81870778 (BR), 81600858 (BR), 81870759 (LC), 81991500 (JYL), 81991501 (JYL); Applied Basic Research Programs of Sichuan Province, 2020YJ0227 (BR); the Youth Grant of the Science and Technology Department of Sichuan Province, China, 2017JQ0028 (LC); Innovative Research Team Program of Sichuan Province (LC); Fund of State Key Laboratory of Oral Diseases (SKLOD201913) (BR).

## Availability of data and materials

Raw sequences of 16S rDNA and transcrptome genes are available in NCBI SRA database (www.ncbi.nlm.nih.gov/sra) under accession number PRJNA640899, PRJNA669544, and PRJNA643633. Raw data of experiments can be accessed through the URL (https://dataview.ncbi.nlm.nih.gov/?search=SUB7647103), (https://dataview.ncbi.nlm.nih.gov/?search=SUB8393147&archive=bioproject), and (https://dataview.ncbi.nlm.nih.gov/?search=SUB7713416&archive=bioproject).

## Ethics approval and consent to participate

Animal ethical statement (AMUWEC2019000) approved by Laboratory Animal Welfare Ethics Committee of the Third Military Medical University.

## Consent for publication

Not applicable.

## Competing interests

The authors declare that they have no competing interests.

## Author details

^1^State Key Laboratory of Oral Diseases & National Clinical Research Center for Oral Diseases & Department of Cariology and Endodontics, West China Hospital of Stomatology, Sichuan University, Chengdu 610044, China. ^2^ Department of Pathogenic Biology, Third Military Medical University, Chongqing 400038, China.

